# *In vivo* clonal analysis reveals spatiotemporal regulation of thalamic nucleogenesis

**DOI:** 10.1101/238964

**Authors:** Samuel Z.H. Wong, Earl Parker Scott, Wenhui Mu, Xize Guo, Ella Borgenheimer, Madeline Freeman, Guo-li Ming, Qing-Feng Wu, Hongjun Song, Yasushi Nakagawa

## Abstract

The thalamus, a crucial regulator of cortical functions, is composed of many nuclei arranged in a spatially complex pattern. Thalamic neurogenesis occurs over a short period during mammalian embryonic development. These features have hampered the effort to understand how regionalization, cell divisions and fate specification are coordinated and produce a wide array of nuclei that exhibit distinct patterns of gene expression and functions. Here, we performed *in vivo* clonal analysis to track the divisions of individual progenitor cells and spatial allocation of their progeny in the developing mouse thalamus. Quantitative analysis of clone compositions revealed evidence for sequential generation of distinct sets of thalamic nuclei based on the location of the founder progenitor cells. Furthermore, we identified intermediate progenitor cells that produced neurons populating more than one thalamic nuclei, indicating a prolonged specification of nuclear fate. Our study reveals an organizational principle that governs the spatial and temporal progression of cell divisions and fate specification, and provides a framework for studying cellular heterogeneity and connectivity in the mammalian thalamus.

## INTRODUCTION

A large portion of the vertebrate brain is organized so that groups of neurons that share similar properties are arranged in aggregates called nuclei. Developmental mechanisms underlying the formation of such nuclear brain structures are much less understood compared with the mechanisms that regulate the formation of laminar structures including the cerebral cortex and retina. The thalamus contains dozens of nuclei that play crucial roles in controlling cortical functions (Jones, 2007; Sherman, 2016). Most thalamic nuclei are populated by excitatory neurons that project axons to the cerebral cortex. Some of these nuclei receive substantial subcortical inputs and convey sensory and motor information to specific areas in the neocortex, whereas other nuclei receive inputs mostly from the cortex and serve as a hub for cortico-cortical communications. Thalamic nuclei can be distinguished by morphology and gene expression as well as by areal and laminar specificity of their axonal projections to the cortex. Genetic fate-mapping at the population level revealed that neurons that populate cortex-projecting nuclei are derived from a progenitor domain marked by the proneural basic helix-loop-helix (bHLH) transcription factors NEUROG1 and NEUROG2 (Fig. 1A, pTH-C domain; (Vue et al., 2007)). In addition to these “excitatory nuclei”, the thalamus also contains “inhibitory nuclei” that are exclusively composed of GABAergic neurons, which include the intergeniculate leaflet (IGL) and ventral lateral geniculate (vLG) nucleus. Notably, recent studies identified a small progenitor domain (pTH-R) that is located rostral to the pTH-C domain during neurogenesis and expresses the proneural bHLH factors ASCL1 and TAL1. Population-level fate-mapping studies using these markers found that pTH-R contributes to IGL and vLG nuclei (Fig. 1A; (Jeong et al., 2011; Vue et al., 2007)). However, due to the complex spatial arrangement of all thalamic nuclei and the narrow time window during which neurons for these nuclei are produced (Angevine, 1970), little is known about how individual progenitor cells undergo divisions and fate specification and eventually produce neurons and glia that populate different thalamic nuclei.

**Figure 1.**
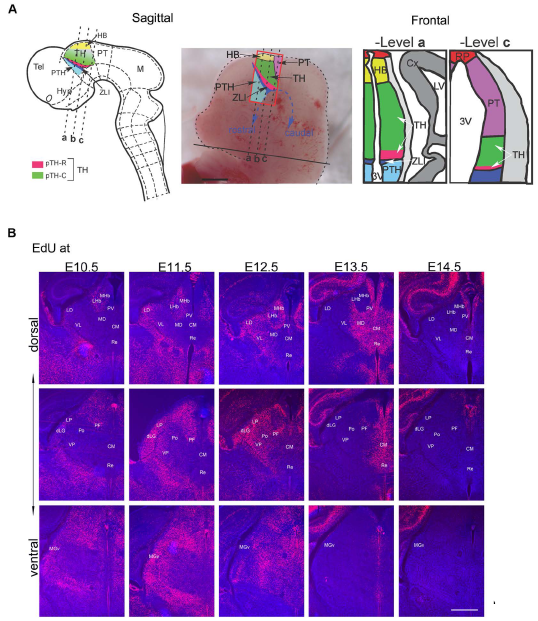
Outside-in temporal specification of thalamic nucleogenesis. (**A**) A schematic of the thalamus (TH) in the caudal diencephalon of the mouse embryo. In the sagittal schematic (left two panels), the embryo is facing left. The thalamus is bordered by the pretectum caudally, the habenula dorsally, the basal plate ventrally, and by the zona limitans intrathalamica (ZLI) rostrally. The three dashed lines represent the planes of sections shown in the frontal schematic (right two panels). In this study, we used Cre drivers of three genes, *Gli1, Olig3* and *Neurog1.* Within the color-labeled regions of the diencephalon, *Gli1* is expressed in pTH-C (rostro-ventral region near the pTH-R domain, shown in darker green) and the prethalamus (light blue) (Vue et al., 2009). *Olig3* is expressed in pTH-C (green) and pTH-R (pink) domains of the thalamus as well as in the zona limitans intrathalamica (dark blue) (Vue et al., 2007). *Neurog1* is expressed in the habenula (yellow), pTH-C (green) and zona limitans intrathalamica (dark blue) (Vue et al., 2007). TH = thalamus, Tel = telencephalon, PTH = prethalamus, Hyp = hypothalamus, ZLI = zonas limitans intrathalamica, HB = habenula, PT = pretectum, M = midbrain, Cx = cortex, RP = roof plate, LV = lateral ventricle, 3V = third ventricle. (**B**) Frontal sections of the forebrain showing EdU birthdating in the neonatal mouse thalamus with EdU injections at E10.5, E11.5, E12.5, E13.5 or E14.5. Three representative levels (dorsal, middle and ventral) are shown for each stage of EdU injection. In all three levels, there is a general trend in which neurons of laterally located nuclei are born before neurons of more medially located nuclei, demonstrating the outside-in pattern of neurogenesis. Scale bar: 500μm.

As demonstrated in invertebrate brains (Kohwi and Doe, 2013), determination of precise cell lineage at the single-cell level is key to understanding the molecular and cellular mechanisms that underlie the formation of complex brain structures like the mammalian thalamus. In this study, we performed clonal analyses of neural progenitor cells of the thalamus using MADM (mosaic analysis with double markers)-based genetic lineage tracing. Quantitative analysis of clone size and distribution obtained with two inducible Cre drivers allowed us to determine the division patterns of individual neuroepithelial and radial glial cells and their differential contributions to distinct thalamic nuclei. Furthermore, we used a third Cre driver to label more differentiated, basally dividing progenitor cells and captured the fate of their progeny generated in the last few cell divisions. These results reveal how patterning, cell division and cell fate specification are coupled in a spatially and temporally specific manner to generate distinct thalamic nuclei during development.

## RESULTS

### Birth-dating of thalamic neurons using a thymidine analog

To investigate the temporal progression of thalamic nucleogenesis in mice, we first performed EdU (ethynyldeoxyuridine) birth-dating by labeling progenitor cells during the S-phase of the cell cycle at embryonic day 9.5 (E9.5), E10.5, E11.5, E12.5, E13.5 and E14.5. We analyzed the distribution of EdU-positive cells at postnatal day 0 (P0) (Fig. 1B, Fig. S1). No labeling was detected in the thalamus with EdU injections at E9.5. With EdU injections at E10.5, robust labeling was detected in ventral medial geniculate (MGv), dorsal lateral geniculate (dLG), lateral posterior (LP) and lateral dorsal (LD) nuclei. With the exception of these earliest-generated nuclei and the latest-generated, paraventricular (PV) nucleus, most thalamic nuclei became post-mitotic between E11.5 and E13.5. The lateral-to-medial neurogenic gradient was evident throughout the thalamus, consistent with previous studies (Altman and Bayer, 1988; Angevine, 1970). Some nuclei (e.g., dLG, LP, LD) were generated over at least 48 hours (E10.5 to E12.5), whereas others (e.g., centromedian; CM, reuniens/rhomboid; Re/Rh, mediodorsal; MD) were generated within one day around E13.5. Thus, thalamic neurogenesis occurs over a short span of embryonic development and likely involves a complex pattern of division and fate specification at the level of individual progenitor cells.

### Establishment of clonal lineage tracing strategy using MADM

In order to determine the temporal and spatial dynamics of neural progenitor cells in the thalamus, we performed genetic lineage tracing at the clonal level by using the mosaic analysis with double markers (MADM) system (Fig. S2A; Hippenmeyer et al., 2010; Zong et al., 2005). When the frequency of Cre-mediated recombination is kept low by titrating the tamoxifen dosage, the sparse Green/Red (G/R) clones generated by “G2-X” type segregation (see Fig. S2 A for definition) would allow us to determine the division pattern and fate of the two daughter cells that are derived from the original progenitor cell undergoing recombination. The yellow (Y) clones produced by “G2-Z” segregation of chromosomes can be defined as hemi-clones because their sister clones are not labeled.

In this study, three inducible Cre driver lines, *Gli1^CreERT2^, Olig3^CreERT2^* and *Neurog1^CreERT2^* were combined with *MADM-11* reporters. Both *Gli1* and *Olig3* are expressed in neuroepithelial and radial glial cells in the thalamus (Vue et al., 2009). *Gli1* is a direct target gene of Sonic Hedgehog signaling pathway and is expressed highly in the rostral part of the thalamus as well as in other regions in the forebrain including the prethalamus and hypothalamus. OLIG3 is expressed in the entire thalamic ventricular zone and in the zona limitans intrathalamica (ZLI), a secondary organizer located immediately rostral to the thalamus (Fig. 1A), whereas its expression is minimal in other forebrain regions (Vue et al., 2007; see Fig. 1A). In the thalamus, NEUROG1 is largely excluded from Notch-ICD (intracellular domain of NOTCH)-expressing neuroepithelial and radial glial cells, and is expressed in basally dividing progenitor cells in the pTH-C domain (Wang et al., 2011).

To validate the recombination patterns of each CreER driver, we performed a population analysis of reporter mice. The *Rosa-H2B-GFP* reporter line crossed with the *Gli1^CreERT2^* driver, and the *Ai6 ZSGreen* reporter crossed with *Olig3^CreERT2^* or *Neurog1^CreERT2^* driver showed labeling of progenitor cells within the thalamic primordium a day after tamoxifen administration (E10.5 for *Gli1,* E11.5 for *Olig3* and E12.5 for *Neurog1).* At E18.5, labeling was detected throughout the thalamus (Fig. 2A).

**Figure 2.**
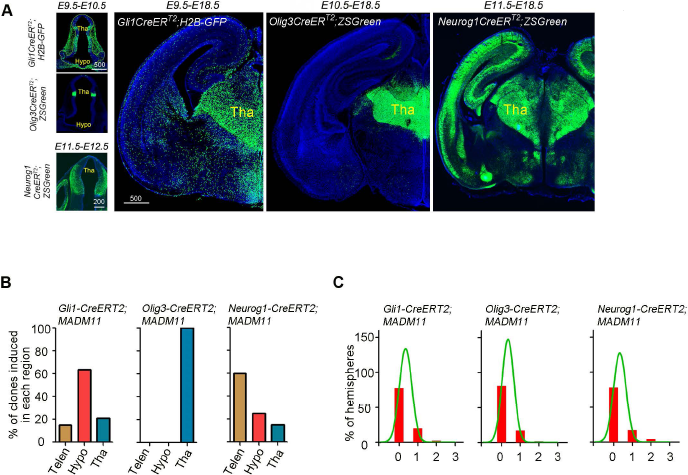
Clonal lineage tracing of thalamic progenitors using MADM. (**A**) Sample images of frontal sections from *Gli1^CreERT2^;H2B-GFP, Olig3^CreERT2^;ZSGreen* and *Neurog1^CreERT2^;ZSGreen* mouse brains counter-stained with DAPI to show labeling of progenitor cells or their progeny. The left three panels show labeling one day after tamoxifen administration (E10.5 for *Gli1^CreERT2^,* E11.5 for *Olig3^CreERT2^* and E12.5 for *Neurog1^CreERT2^).* Tha; thalamus, Hypo; hypothalamus. Scale bar: 500μm. (**B**) Bar graphs showing regional distribution of labeled clones in the forebrain of *Gli1^CreERT2^, Olig3^CreERT2^ and Neurog1^CreERT2^* mouse lines crossed with *MADM11* reporter mice. Brains were analyzed one day after tamoxifen administration. Telen, telencephalon; Tha, thalamus; Hypo, hypothalamus. (**C**) Bar graphs showing frequencies of thalamic clones (green/red or yellow) per hemisphere labeled one day after tamoxifen administration for the *Gli1^CreERT2^;MADM11, Olig3^CreERT2^;MADM11* and *Neurog1^CreERT2^;MADM11* mice. More than 75% of hemispheres had no recombined clones. The curves represent the best non-linear fit.

We then crossed each CreER driver with *MADM-11^GT/GT^* mice, and the *CreER^+/-^; MADM-11^GT/GT^* mice were crossed with *MADM-11^TG/TG^* mice. A day after the tamoxifen administration, MADM brains for all three CreER drivers had G/R and Y clones (see Fig. S2 A for definition) in thalamic primordium (Fig. 2B), and the apical and basal processes of the labeled cells in *Gli1^CreERT2^* MADM brains were positive for Nestin, a neuroepithelial and radial glial cell marker (Fig. S2B). Many clones were produced in the thalamus of *Gli1^CreERT2^* MADM brains with tamoxifen administration at E9.5. In contrast, *Olig3^CreERT2^* MADM clones were obtained only after tamoxifen administration at E10.5 or later. This is likely due to the difference in the onset of the expression of the Cre drivers. In addition, expression of *Gli1* has a rostral-high, caudal-low gradient within the thalamus (Vue et al., 2009) and *Gli1* MADM clones tended to be concentrated in the rostral part of the thalamus. Thus, using both *Gli1* and *Olig3* CreER drivers would allow us to fully trace the lineage of spatially and temporally divergent neuroepithelial and radial glial cell populations in the thalamus. In addition, the *Neurog1* CreER driver would allow labeling of progenitor cells at a later stage of neurogenesis. Both *Gli1* and *Neurog1* CreER drivers labeled progenitor cells widely in the forebrain, whereas the *Olig3* driver was specific to the thalamus (Fig. 2B). By adjusting the dosage of tamoxifen for each CreER driver, we achieved a low probability of labeling a pair of progeny generated by a single recombined progenitor cell in the thalamus (Fig. 2C). The percentage of hemispheres that had at least one G/R or Y clone a day after the Cre activation was less than 25% (~22% for *Gli1^CreER^,* ~19% for *Olig3^CreER^* and ~22% for *Neurog1^CreER^),* and almost no hemispheres had two clones (~2% for *Gli1^CreER^,* ~3% *Olig3^CreER^* and ~5% for *Neurog1^CreER^),* which validated our methodology for clonal analysis of thalamic progenitor cells. For the *Olig3^CreER^* driver, we analyzed 186 hemispheres (27 litters) of *CreER+* E18.5 mice and 112 hemispheres (23 litters) of *CreER+* P21 mice. Out of these brains, we detected 72 and 22 G/R clones, respectively. For the *Gli1^CreER^* driver, we analyzed 194 hemispheres (37 litters) of *CreER+* E18.5 mice and 112 hemispheres (29 litters) of *CreER+* P21 mice. Out of these brains, we found 27 and 24 G/R clones, respectively. For the *Neurog1^CreER^* driver, we analyzed 83 hemispheres (17 litters) of *CreER+* E18.5 mice and found 55 G/R clones. In order to avoid analyzing potentially mixed clones that arise from two or more recombined cells, we excluded *Olig3^CreERT2^* E18.5 brains from two litters in which every hemisphere had fluorescently labeled cells in the thalamus.

### Analysis of clonal division patterns of thalamic progenitor cells

A major advantage of using the MADM system for lineage tracing is that it allows for analysis of the division pattern of the recombined single progenitor cell based on the numbers of green cells and red cells in each G/R clone. In early embryonic neocortex, neuroepithelial cells undergo symmetric proliferative divisions to increase the pool size of progenitor cells. As neurogenesis starts, progenitor cells divide asymmetrically and produce a self-renewed radial glial cell and a more differentiated cell, either a post-mitotic neuron or an intermediate progenitor cell (IPC). IPCs in embryonic neocortex express the T-box transcription factor TBR2 and divide symmetrically in basal locations away from the surface of the lateral ventricle (Kriegstein and Alvarez-Buylla, 2009; Florio and Huttner, 2014; Mihalas et al., 2016; Yao et al., 2016). Although TBR2 is not expressed in embryonic thalamus, basally dividing cells do exist that express the proneural basic helix-loop-helix transcription factors NEUROG1 and NEUROG2 (Wang et al., 2011). Since neocortical IPCs also express NEUROG2 (Miyata et al., 2001), we hypothesized that Neurogenin-expressing, basally dividing cells in the thalamus are IPCs. In E18.5 *Neurog1^CreERT2^* MADM brains, 78% of G/R clones had 4 or fewer cells (Fig. 3A) and all labeled cells had a neuronal morphology. Thus, a majority of Neurog1-expressing thalamic progenitor cells are intermediate neuronal progenitor cells with a limited proliferative capacity.

**Figure 3.**
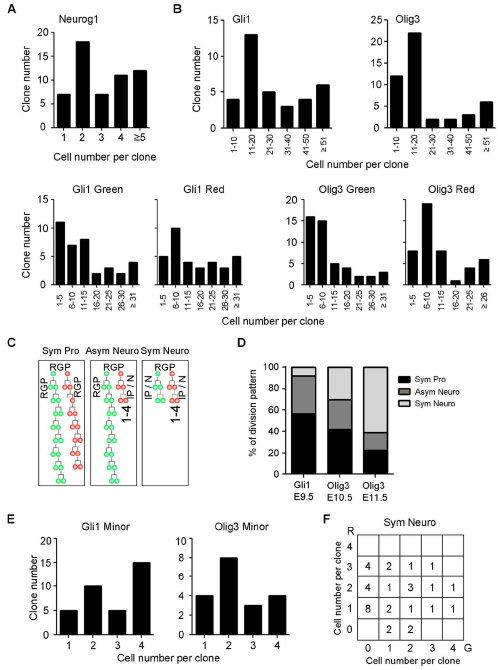
Temporal change of cell divisions during thalamic nucleogenesis. (**A**) Bar graph showing a breakdown of the cell number per clone for all *Neurog1^CreERT2^* labeled clones. (**B**) Top: Bar graphs showing breakdown of the cell number per clone for all *Gli1^CreERT2^* and *Olig3^CreERT2^* labeled clones. Bottom: Bar graphs showing breakdown of the cell number per clone for all *Gli1^CreERT2^* and *Olig3^CreERT2^* labeled green/red hemi-clones. (**C**) A schematic depicting the three defined types of clonal division patterns: symmetric proliferative (“Sym Pro”), asymmetric neurogenic (“Asym Neuro”) and other (“Sym Neuro”). (**D**) Bar graph showing the percentage of division pattern within clones labelled using *Gli1* and *Olig3* driver lines with tamoxifen administration at E9.5 for *Gli1* and E10.5/E11.5 for *Olig3.* (**E**) Bar graph showing the distribution of the size of minority hemi-clones in asymmetric neurogenic clones of *Gli1* and *Olig3* MADM brains. (**F**) Graph showing the distribution of the numbers of green (G) and red (R) cells of symmetric neurogenic clones of *Gli1* and *Olig3* MADM brains.

In contrast to the small size of *Neurog1* MADM clones, most clones in *Gli1* and *Olig3* MADM brains had more than 10 cells (Fig. 3B). Large G/R clones with more than 4 cells in both green and red progeny are likely to originate from a recombined progenitor cell that produced two radial progenitors, not *Neurog1-* expressing progenitor cells. We defined these clones as symmetric proliferative (Fig. 3C; “Sym Pro”). G/R clones in which either green or red hemi-clone contained 4 or fewer cells were defined as asymmetric self-renewing/neurogenic, which likely produced one radial progenitor and an IPC, or one radial progenitor and a postmitotic neuron (Fig. 3C, “Asym Neuro”). G/R clones with 4 or fewer cells in both hemi-clones were defined as symmetric neurogenic (Fig. 3C, “Sym Neuro”).

Both *Gli1* and *Olig3* clones exhibited a combination of the three division patterns described in Fig. 3C (Fig. 3D). When recombination was induced in *Gli1^CreER^* mice at E9.5, ~57% of the clones were of the symmetric proliferative type, indicating that a majority of recombined progenitor cells at E9.5 were neuroepithelial cells undergoing symmetric expansions. In *Olig3^CreER^* mice, this fraction was ~42% at E10.5 and ~22% at E11.5. In addition, at E11.5, ~61% of the clones were of the symmetric neurogenic type. The average clone size was smaller for a later Cre activation (22.5 for E10.5 and 9.5 for E11.5). The size of the minority hemi-clones in asymmetric clones (Fig. 3E) and the composition of symmetric neurogenic clones (Fig. 3F) both showed a large variation. These results together indicate a transition of cell division mode in the thalamus from symmetric proliferative to asymmetric to symmetric neurogenic, similar to the neocortex (Gao et al., 2014).

### Spatial organization of clonal units reveals principles of thalamic nucleogenesis

To investigate the spatial organization of clonal units during thalamic nucleogenesis, we performed three-dimensional (3D) reconstructions of serial brain sections containing all labeled cells (Fig. 4A and Fig. S3) (Bonaguidi et al. 2011) and assigned each labeled cell to a specific thalamic nucleus using a published atlas (Paxinos, 2006), data from our previous study (Nakagawa and O'Leary, 2001), and a newly generated custom atlas (Fig. S4). Asymmetric G/R clones generated 11.9 ± 0.87 cells (Fig. 4B) that populated 3.3 ± 0.26 nuclei (Fig. 4C). This demonstrates that each thalamic radial glial cell produces neurons that populate multiple nuclei after asymmetric divisions. The average number of nuclei populated by progeny of asymmetrically dividing progenitor cells was significantly smaller than the number for symmetric proliferative clones (Fig. 4C), suggesting that the transition from symmetric proliferative to asymmetric division restricts the nuclear fate potential. Within asymmetric clones, “minority” hemi-clones (i.e. G or R hemi-clones containing 4 or fewer cells) exhibited significantly more fate restriction than “majority” hemi-clones based on the average number of populated nuclei (Fig. 4C). However, some minority hemi-clones still populated more than one nuclei, implying that the fate of thalamic nuclei may not be fully restricted even in terminally dividing cells. The three principal sensory nuclei (VP, dLG, MGv) are among the earliest-born in the thalamus (Fig. 1B). Approximately 61% of majority hemi-clones were composed of both principal sensory nuclei and other later-born nuclei, whereas only 12% of them had cells only in principal sensory nuclei. Strikingly, 44% of minority hemi-clones showed an exclusive contribution to principal sensory nuclei (Fig. 4D). This suggests that thalamic radial glial cells sequentially generate neurons that populate different nuclei as they undergo asymmetric divisions. This pattern is reminiscent of the sequential generation of deep- and upper-layer neurons from individual radial glial cells in the embryonic neocortex (Gao et al., 2014).

**Figure 4.**
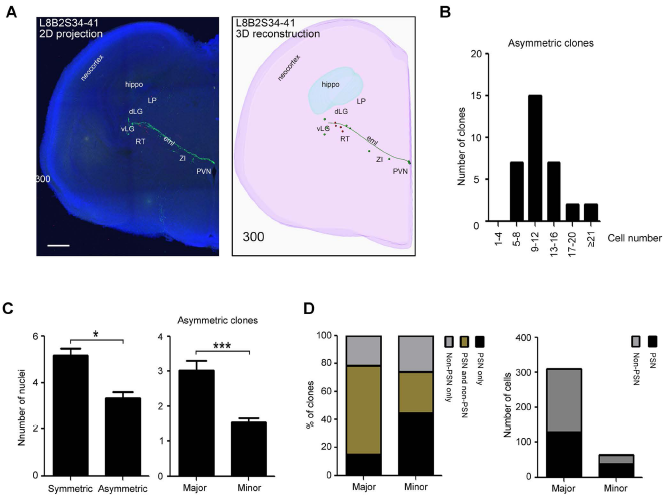
Cell division patterns reveal the fate specification of early-born neurons in thalamus. (**A**) Brains were reconstructed from serial sections cumulatively containing all labelled cells within a single clone using IMARIS. Shown is a sample representative 2D projection (left panel) and 3D reconstruction (right panel) of thalamic clone derived from a *Gli1^CreERT2^* MADM E18.5 brain. Violescent stereostructure sketches the contours of the hemisphere, the green filament shows the process of a residual radial glial cell, and green and red spots represent labelled GFP+ and tdTomato+ cells, respectively. Scale bar: 300μm. (**B**) Bar graph showing the distribution of the size (cell number) of asymmetric clones. (**C**) Left: bar graph showing the average number of nuclei populated in symmetric and asymmetric G/R clones. Symmetric proliferative clones contributed to more thalamic nuclei than asymmetric neurogenic clones. Right: bar graph showing the average number of nuclei populated in majority (“major”) and minority (“minor”) hemi-clones within asymmetric clones. (**D**) Bar graph showing the percentage of clones (left panel) and number of cells (right panel) populating principal sensory nuclei (PSN) in majority (“major”) and minority (“minor”) hemi-clones within asymmetric clones.

### Individual thalamic clones exhibit distinct patterns of nuclear contributions

We next tested if individual radial glial clones contribute to specific subsets of thalamic nuclei. To analyze our results computationally, we generated a data matrix in which rows and columns represented individual G/R clones and thalamic nuclei, respectively. The numeric value in each cell of the matrix was computed (see Materials and Methods) so that it indicated the frequency of progeny in a given clone that was located in each thalamic nucleus. This configuration is analogous to the matrix used for single-cell RNA-seq analysis, in which rows, columns and values represent individual cells, genes and expression levels, respectively. We then applied principal component analysis or unsupervised hierarchical clustering analysis for classification of thalamic MADM clones.

Principal component analysis of asymmetric G/R clones identified five clusters that show distinct patterns of nuclear contribution (Fig. 5A). Three of the five clusters had major contributions to principal sensory nuclei (yellow, green and red clusters as shown in Fig. 5A). Of these, the yellow cluster contained clones that contributed to principal sensory nuclei and other nuclei including ventromedial (VM) and reuniens/rhomboid (Re/Rh). Clones in the green cluster included cells in principal sensory nuclei and medial ventral field (Fig. S4C; see Materials and Methods for definition). VM and Re/Rh and the medial ventral field are all located medial to principal sensory nuclei, and neurons in these nuclei are born later than those in principal sensory nuclei (Fig. 1B). Importantly, in seven out of eight green clones, the minority hemi-clones contributed exclusively to principal sensory nuclei. This result suggests that asymmetrically dividing thalamic radial progenitor cells first generate early-born, principal sensory nuclei and then late-born, medially located nuclei. It also suggests a strong lineage relationship among the three principal sensory nuclei, which share many properties including laminar and areal patterns of axon projections to primary sensory areas of the neocortex (Clasca et al., 2012; Jones, 2007), gene expression (Frangeul et al., 2016; Nagalski et al., 2016) and patterns of propagating spontaneous activity during embryogenesis (Moreno-Juan et al., 2017).

**Figure 5.**
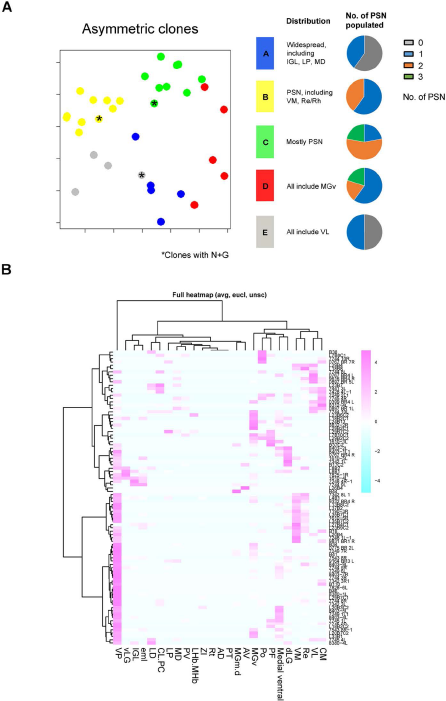
Spatial specification of thalamic nucleogenesis. (**A**) Principal component analysis of asymmetric G/R clones in *Olig3* and *Gli1* MADM brains at both E18.5 and P21, together with a summary of nuclei populated in each cluster. Asterisks label clones that contain both neurons and astrocytes. (**B**) Unsupervised hierarchical clustering analysis of all types of G/R clones including symmetric proliferative clones. The heatmap shows the distance between different thalamic nuclei.

The other two (blue and grey) clusters included clones that have few cells in principal sensory nuclei (Fig. 5A). All the clones in the grey cluster contributed to the ventrolateral (VL) nucleus. The remaining nuclei represented in this cluster were diverse and included centromedian/intermediodorsal (CM/IMD), mediodorsal (MD) and Re/Rh. Many of these nuclei were located more dorsally and caudally within the thalamus compared with principal sensory nuclei, demonstrating that nuclei that are close to each other in physical location are likely to share the same lineage. This result provides single-cell based evidence consistent with our previous population-based study, in which gradients of gene expression within the pTH-C domain corresponded to preferential generation of progeny along the rostro-ventral to caudo-dorsal axis of the thalamus (Vue et al., 2007).

The blue cluster contained clones that were relatively more diverse. Here, two clones contained cells in intergeniculate leaflet (IGL) and either MGv or VP. The IGL nucleus receives axonal projections from non-image forming retinal ganglion cells and are known to be derived from the pTH-R domain at the population level (Delogu et al., 2012; Jager et al., 2016; Jeong et al., 2011; Vue et al., 2007). The surprising co-existence of pTH-C derived cells (in principal sensory nuclei) and pTH-R derived cells (in IGL) in the same clones suggests that during early stages, some progenitor cells might still be uncommitted to either the pTH-C or the pTH-R fate.

The existence of clone clusters suggests that some thalamic nuclei are more likely to share the cell lineage than other nuclei. In order to assess the clustering of clones and the “lineage distance” between nuclei, we performed unsupervised hierarchical clustering analysis and generated a heatmap. Because a majority of the clones that contributed to late-born nuclei were of the symmetric proliferative type, we included both asymmetric and symmetric G/R clones in this analysis in order to cluster the entire set of thalamic nuclei. We found that certain groups of nuclei were more likely to originate from common progenitor cells at E18.5 (Fig. 5B). Examples of additional clusters of nuclei identified in this analysis include 1) VM, VL, CM/IMD and Re/Rh, 2) LD and CL/PC, 3) LP, MD and PV, 4) MGv, dLG, Po, medial ventral field and PF, and 5) vLG and IGL. These clusters of nuclei are generally arranged radially and are located at similar rostro-caudal and dorso-ventral positions within the thalamus. The presence of lineage-based clusters of nuclei shown here suggests that the distribution of progeny generated from thalamic progenitors appears to follow distinct nuclei-specific patterns.

### Position of radial progenitor cells dictates domain-specific distribution of clones

To determine if the clustering of clones based on cell lineage reflects the distinct positions of starting progenitor cells within the embryonic thalamus, we took advantage of the finding that some clones, especially the ones labeled at E9.5 in the *Gli1^CreER^* line, retained a cell at the ventricular surface with a long radial process spanning the entire thalamic wall.

In order to use the locations of these cells as a proxy for the position of the founder progenitor cell at early embryonic stages, we defined four domains of these radial glial cells within the E18.5 thalamus (Fig. 6A). “Dorsal-rostral (DR)” corresponded to the most rostral domain in relatively dorsal sections, whereas the “dorsal-middle (DM)” and “dorsal-caudal (DC)” domains were located progressively more caudally. The “ventral (V)” corresponded to the domain in more ventral sections near the basal plate. Two-dimensional shapes of clones derived from each of the four domains are shown in Fig. 6B. A hierarchical clustering analysis revealed that clones containing radial glial cells in the DR and DM domains heavily contributed to principal sensory nuclei (Fig. 6C); many of these clones contained cells in both principal sensory nuclei and more medially located non-sensory nuclei including VM, Re/Rh and the medial ventral field. Some DR clones contained cells in both “excitatory nuclei” composed of cortex-projecting neurons and “inhibitory nuclei” containing GABAergic neurons. Thus, DR and DM clones appear to correspond to the yellow, green and red clones in Fig. 5A. DC and V clones had a large contribution to dorsally located, non-sensory nuclei and a small contribution to principal sensory nuclei except MGv. These results suggest that thalamic nuclei that are related in ontogeny originate from similar locations within the embryonic thalamus and support the model wherein individual thalamic radial glial cells form a radial column of progeny that spans a defined subset of thalamic nuclei.

**Figure 6.**
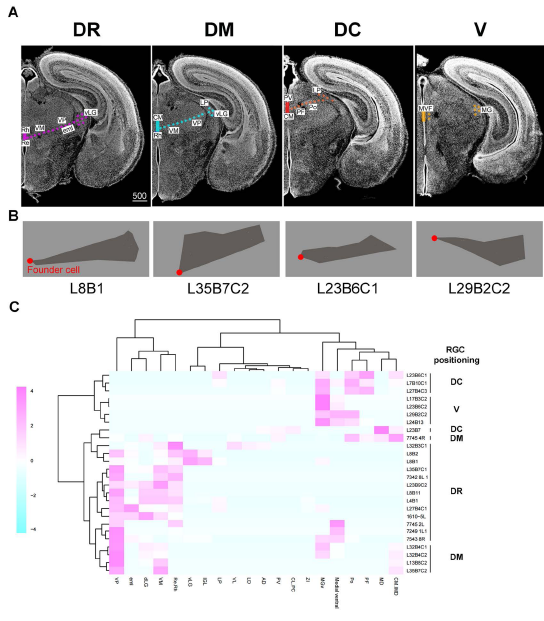
RGC position dictates domain-specific distribution of clones. (**A**) The four micrographs show frontal sections of E18.5 forebrain, in which colored solid bars indicate the four zones, DR (dorsal rostral), DM (dorsal middle), DC (dorsal caudal) and V (ventral), of GC positions. Scale bar: 500μm. Bottom: four faying surface plots illustrate representative shapes of the respective clones. (**B**) 2-D shape of a representative clone for each location type shown in Fig. 6A. The red dot indicates the position of the residual radial glial cell. (**C**) Clustering analysis of clones based on the position of radial glial cells (RGCs).

### Analysis of glia-containing clones reveals rare segregation of neurons and glia across nuclei

Out of the 46 G/R P21 clones obtained from *Gli1^CreER^* and *Olig3^CreER^* drivers, 17 clones were composed of both neurons and glia. Clones containing glia were distributed in multiple PCA clusters (glia-containing clones in asymmetric G/R clones are indicated by an asterisk in Fig. 5A), suggesting that the ontogeny of thalamic glial cells is heterogeneous and that gliogenesis is not restricted to specific thalamic progenitor cell populations.

Many postnatal MADM brains contained glial cells. The average number of glial cells was 24.7 ±11.4 cells in the P21 glia-containing clones. Although the number of glial cells did not correlate with the number of neurons in the P21 clones, our data suggested a significant linear correlation between the number of glia and total cells (Fig. S5A). Among all of the postnatal clones, three were composed mostly of glial cells, suggesting the existence of glial lineage-specific radial glial precursors. The distribution analysis of glial number in the P21 clones showed that the three clones with a large number of glia were outliers (Fig. S5B). After removing these clones, we did not find any correlation between the glial number and the neuronal number or between glial number and total cell number in glia-containing clones (Fig. S5C). The *Gli1* clones contained more glia than *Olig3* clones (Fig. S5D), which might be due to the earlier labelling of radial glial precursor in the *Gli1* clones (at E9.5) than in *Olig3* clones (E10.5 or E11.5). Bipotent radial glial progenitors producing neurons and astrocytes (N+A) generated more neurons than unipotent progenitors producing neurons only (Fig. S4E). We did not collect sufficient clones derived from bipotent progenitors producing neurons and oligodendrocytes (N+O) and tripotent progenitors producing all three types of neural cells (N+A+O) to compare with unipotent progenitors.

The majority of symmetric proliferative and asymmetric neurogenic P21 clones contained both neurons and glia (Fig. S5F). Together, our data suggest that a majority of radial glial cells in different progenitor domains are multipotent and generate glial cells by P21.

### Clonal lineage tracing of basal progenitor cells shows prolonged specification of thalamic nuclear fate

Analysis of asymmetric clones of *Gli1^CreER^* and *Olig3^CreER^* MADM brains revealed minority hemi-clones that included cells in more than one thalamic nuclei (Fig. 4C). This suggested that progenitor cells undergoing their last few rounds of divisions are still not yet specified to the fate of a single nucleus. As described above (Fig. 3A), a large majority of *Neurog1^CreER^* MADM clones contained 4 or fewer cells with a neuronal morphology, indicating that *Neurog1*-expressing thalamic progenitor cells are IPCs. Taking further advantage of the MADM system, we analyzed the division patterns and fates of these cells.

At the population level, timed activation of Cre recombinase at E11.5 resulted in expression of the ZSGreen reporter in the entire thalamus except the medial-most region including PV, CM and Re/Rh nuclei (Fig. 2A, 7A). These medial nuclei began to be labeled by ZSGreen with the Cre activation at E12.5 (Fig. 7A). This is one day before they become robustly labeled through EdU birth-dating (Fig. 1B). Similarly, VL starts to be labeled by ZSGreen at E11.5 and by EdU at E12.5, indicating that at least for the above thalamic nuclei, progenitor cells express *Neurog1* before they undergo the final division.

**Figure 7.**
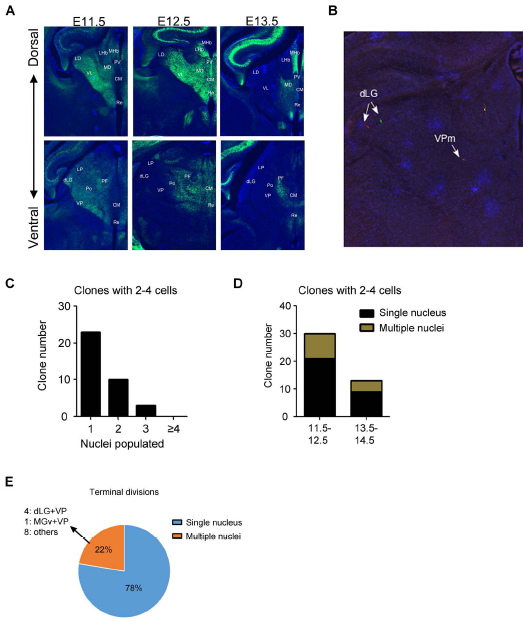
Clonal lineage tracing of basal progenitor cells shows prolonged nuclear specification of thalamic cells. (**A**) Left: sample images of frontal sections from E18.5 *Neurog1^CreERT2^;ZSGreen* mouse brains to show labeling of progeny of *Neurog1*-expressing progenitor cells when tamoxifen is administered at E11.5, E12.5 or E13.5. Sections at three different dorso-ventral levels are shown for each stage of tamoxifen administration. (**B**) Sample image of a *Neurog1^CreERT2^* clone at E18.5. The colonization of two nuclei within a single clone is shown. Scale bar: 300μm. (**C**) Bar graph showing the number of nuclei populated for *Neurog1^CreERT2^* labeled clones with 2 to 4 cells. (**D**) Bar graph showing the percentage of clones populating single or multiple nuclei for *Neurog1-*labeled clones with 2-4 cells. The left bar indicates clones that are activated at E11.5 or E12.5, and the right bar indicates clones activated at E13.5 or E14.5. (**E**) Pie chart showing the percentage of clones populating single or multiple nuclei for clones undergoing terminal divisions. The breakdown of nuclei populated for clones colonizing multiple nuclei is shown.

We next performed clonal lineage tracing with MADM using the *Neurog1* CreER driver. Activation of the Cre recombinase at E11.5, E12.5, E13.5 or E14.5 followed by analysis at E18.5 revealed that a vast majority of MADM clones contained 4 or fewer cells (43 out of 55 G/R clones; Fig. 3A). Surprisingly, many of these clones (13 out of 36 G/R clones with 2-4 cells) contained neurons that populated more than one thalamic nuclei (Fig. 7B,C). Clones covering multiple nuclei were abundant with E11.5 or E12.5 activation (9 out of 30 clones), but there were some with the activation at E13.5 or E14.5 (4 out of 13 clones; Fig. 7D). Four of the multi-nuclei clones spanned two or all three principal sensory nuclei, while others included late-born nuclei like PV and Re/Rh. By taking advantage of the dual-color labeling of the progeny in the MADM system, we also captured 58 terminal divisions for which we could determine whether the last division generated cells in the same nucleus or two different nuclei. Among these 58 final divisions, 13 generated cells in two different nuclei, of which 4 were in dLG and VP and one was in MGv and VP (Fig. 7E), consistent with the ontogenic proximity of these nuclei revealed in the analysis of *Gli1^CreER^* and *Olig3^CreER^* MADM samples (Fig. 5). For individual nuclei, we found that 4 out of 8 terminal divisions that produced a neuron in dLG also produced a neuron in another nucleus (all in VP). Out of 13 terminal divisions that produced a neuron in VP, 7 produced a neuron in another nucleus (4 in dLG, 1 in MGv, 1 in LP, 1 in CL/PC). Thus, dLG and VP nuclei are particularly close to each other in cell lineage and their fates may not diverge at least until the final progenitor cell division. In contrast, only 2 out of 9 terminal divisions that produced a neuron in MGv also produced a neuron in another nucleus (1 each in VP and LP), suggesting that the MGv lineage is largely specified by the time of the final division of *Neurog1-*expressing IPCs.

Taken together, analysis of *Neurog1^CreER^* clones demonstrated that the thalamus contains IPCs and that many of these cells are not yet specified for the fate of a specific nucleus. Strikingly, even the last division of such progenitor cells can generate cells in two distinct nuclei, indicating prolonged mechanisms of nuclear fate specification in developing thalamus.

## DISCUSSION

In this study, we tracked the lineage of thalamic progenitor cells at the clonal level by using the MADM system with three different inducible Cre drivers. The dualcolor labeling of progeny derived from individual progenitors allowed us to examine the temporal progression of progenitor cell differentiation based on division patterns. Our results revealed several cardinal features of thalamic nucleogenesis; 1) contribution to multiple nuclei by individual radial glial cells, 2) temporal progression of nucleogenesis within the lineage, 3) relationship between the position of radial glial cells and the contribution to specific sets of nuclei, and 4) presence of neuronal IPCs and prolonged specification of nuclear fate. We propose that these features are fundamental to the organizing principles of complex brain structures like the thalamus.

### Individual radial glial cells in the thalamus produce many neurons that populate multiple nuclei

Thalamic neurogenesis occurs over a short period of time during embryonic development. In mice, neurons in the earliest-born thalamic nuclei, including dLG and MGv, are first generated at around E10.5, while later-born nuclei are born by E13.5 with an overall “outside-in” pattern (Angevine, 1970; Fig. 1B). Due to the short time span of thalamic neurogenesis and the spatially complex arrangement of dozens of nuclei, it was previously unknown whether each neural progenitor cell produces a limited number of progeny of a single nuclear identity or sequentially generates a large number of neurons that populate multiple nuclei with different birthdates. With the MADM approach, we defined clones that originated from an asymmetrically dividing radial glial cell, and found that each asymmetric clone contains an average of 11.9 cells that populated 3.3 thalamic nuclei. This result provides a general principle of thalamic nucleogenesis in which a single radial glial cell produces many neurons that populate multiple nuclei. By distinguishing the differentially colored hemi-clones derived either from a self-renewed radial glia or a differentiated daughter cell, we found that many radial glial cells labeled at E9.5, E10.5 or E11.5 first produced neurons that populate early-born, principal sensory nuclei including dLG, VP and MGv. Then, in many clones, the same lineage produced neurons that occupy more medially located, later-born nuclei such as Re/Rh and VM (Fig. 1B). A recent study by Gao et al. (2014) analyzed neurogenesis in the neocortex using the MADM approach and found that each asymmetrically dividing radial glia sequentially produces an average of 8.4 cells that populated multiple layers in an inside-out fashion. This indicates that despite the difference between nuclear versus laminar organizations of these two structures, the thalamus and neocortex might share a basic strategy in cell division and differentiation.

### Regional heterogeneity of thalamic progenitor cells in nuclear contributions and neurogenesis

In order to elucidate the heterogeneity among individual cell lineages, we performed principal component analysis of asymmetric MADM clones obtained with *Gli1^CreER^* and *Olig3^CreER^* drivers. This revealed several clusters of clones each of which contributed to a shared set of thalamic nuclei (Fig. 5A). Out of the five clusters identified, three showed significant contributions to principal sensory nuclei, dLG, VP and MGv. One of these clusters (yellow) was composed of principal sensory nuclei and later-born nuclei that lie medial to principal sensory nuclei, indicating that some radial glia are capable of sequentially generating cells that contribute to different sets of nuclei located along the radial axis. The green cluster included few nuclei other than the principal sensory nuclei. Thus, some radial glial lineages may be depleted once the early-born sensory nuclei are generated. The third such cluster (red) had a strong contribution to MGv, the principal auditory nucleus. These clones contributed to a different set of late-born nuclei including PF and Po from the clones of green and yellow clusters. The remaining two clusters of clones had more contributions to caudo-dorsally located nuclei than principal sensory nuclei. Thus, although the principal sensory nuclei generally shared a progenitor cell lineage, the actual pattern of nuclear contribution differed between clones. Clustering of nuclei based on contributions by both symmetric and asymmetric G/R clones revealed additional nuclear groups that are located more caudo-dorsally than principal sensory nuclei (Fig. 5B).

Recently, Shi et al. (2017) reported MADM-based lineage tracing in the postnatal mouse thalamus using the *NestinCreER* driver. They used an unsupervised hierarchical clustering method to cluster both MADM clones and thalamic nuclei based on the distribution of the cells of each clone in different nuclei. The patterns of clone clusters are generally similar to ours (Fig. 5A,B). They found the three principal sensory nuclei and the motor nuclei VA/VL are ontogenetically segregated from three other clusters (MD/Po/LP/CM etc., VM/Re, and AM/AV/PV/LD, etc.). Sherman and colleagues (Sherman et al, 2016; Sherman and Guillery, 2013) have proposed that the three principal sensory and VA/VL nuclei in the thalamus receive strong subcortical input and relay the information to primary sensory and motor areas (“first-order” nuclei). Therefore, Shi et al. concluded that the first-order and higher-order nuclei, which receive most driver input from the cortex, are ontogenetically segregated. Our study found clusters of nuclei (Fig. 5B) that are similar to those detected by Shi et al.. However, there were some differences in the results as well as interpretation. First, our analysis did not find a strong clustering of VL with principal sensory nuclei. Instead, VL more closely segregated with CM, Re and VM (Fig. 5A,B), indicating that the ontogenetic relationship of thalamic nuclei does not necessarily follow the distinction between the first-order versus higher-order nuclei. In addition, another first-order nucleus, anterior nuclear complex, particularly the anterodorsal (AD) nucleus, which receives subcortical driver input from the mammillary body (Sherman and Guillery, 2013, Sherman 2016, Petrof and Sherman, 2009), was ontogenetically distant from the principal sensory nuclei both in our study and in Shi et al., (2017). Thus, we propose that the ontogenetic relationship between thalamic nuclei are defined primarily by their rostro-caudal and dorso-ventral positions instead of functions. Patterns of gene expression (Frangeul et al., 2016; Nagalski et al., 2016; Nakagawa and O’Leary, 2001; Suzuki-Hirano, 2011), laminar specificity of efferent projections to the cortex (Clasca et al., 2012) and sources and types of afferent projections (Sherman, 2016; Sherman and Guillery, 2013) are likely depend on both cell lineage and post-mitotic, potentially extrinsic, mechanisms.

In addition to simply clustering clones and nuclei, we attempted to correlate the locations of original progenitor cells with their nuclear contributions by analyzing clones that retained radial glial cells at E18.5 (Fig. 6). This analysis revealed that the location of the original radial glia predicts the set of nuclei it populates. For example, rostrally located clones (shown as “DR” and “DM” clones in Fig. 6A) strongly contributed to principal sensory nuclei, whereas more caudally located, DC clones contributed to caudo-dorsal nuclei including Po, PF, MD and CM/IMD. Interestingly, some of the most rostrally located (DR) clones included both principal sensory nuclei and IGL/vLG, the two nuclei that are mostly populated by inhibitory neurons. This was unexpected because previous population-based lineage studies found that IGL and vLG nuclei originate from a distinct progenitor domain, pTH-R, not pTH-C that contributes to principal sensory nuclei (Delogu et al., 2012; Jeong et al., 2011; Vue et al., 2007; Vue et al., 2009). Although we did not determine if the neurons in pTH-C-derived principal sensory nuclei that shared the lineage with those in IGL/vLG are glutamatergic or GABAergic, there are almost no GABAergic neurons in VP and MGv in mice and those in dLG are derived from the midbrain and appear only postnatally (Jager et al., 2016). In addition, we observed this type of clone not only at E18.5, but also at P21, making it unlikely that all the neurons in the pTH-C domains in clones of this cluster at E18.5 were still on route to their final destinations in pTH-R derived nuclei. Thus, it is likely that some clones do contribute to both glutamatergic pTH-C nuclei and GABAergic pTH-R nuclei. One possible mechanism for this is that some early progenitors are still uncommitted for either the pTH-C or pTH-R fate. This is consistent with the fact that the pTH-R domain emerges only after neurogenesis starts in the thalamus (Vue et al., 2009).

The location of the originating radial glia were not only correlated with the set of nuclei it populates, but also the temporal patterns of cell division. Most of the clones that contained caudo-dorsally located, late-born nuclei such as mediodorsal (MD), centromedian (CM) or paraventricular (PV) nuclei were found to be of the symmetric proliferative type (26 out of 32 clones containing cells in these nuclei were of symmetric type), demonstrating that upon early Cre activation, progenitor cells that produce neurons of these nuclei are still expanding the pool size by symmetric proliferative divisions (Fig. S6A). Analysis of clones containing a residual radial glia (Fig. 6) indicated that cells of MD, CM and PV nuclei are derived from caudo-dorsally located progenitor cells. In contrast, progenitor cells in the rostral part contributed to principal sensory nuclei, many of which underwent an asymmetric division immediately after the recombination at E9.5 to E11.5 (28 out of 77 clones containing cells in these nuclei were of asymmetric type) (Fig. S6A). Our results indicate that the temporal pattern of transition from symmetric to asymmetric divisions differs among progenitor domains within the thalamus, where the transition occurs earlier in rostro-ventrally located progenitor cells than in the caudo-dorsal cells and produces a cohort of earlier-born nuclei.

### Prolonged specification of nuclear fates in the thalamus

Our previous study showed that similar to the neocortex, the thalamus has a large population of progenitor cells dividing basally away from the surface of the third ventricle (Wang et al., 2011). However, the division patterns and the fate of these cells had remained unknown. By using the *Neurog1^CreER^* driver for clonal lineage analysis of these basal progenitor cells in the thalamus, we found that most of the labeled clones contained four or fewer cells of neuronal morphology. This demonstrates that *Neurog1*-expressing basal progenitor cells in the thalamus are neuron-generating IPCs. The existence of IPCs fits the model of sequential neurogenesis in the thalamus, in which each radial glial cell divides a limited number of times during a short neurogenic period and generates IPCs, and that these IPCs divide once or twice to expand the neuronal output. In the MADM study by Gao et al., (2014), the average size of asymmetric clones in the embryonic neocortex was 8.4 when they were defined as those with 3 or fewer G or R cells. Using the same criteria, the number for the embryonic thalamus in the current study was 11.4, which indicates that the asymmetrically dividing radial glia in the thalamus produce more progeny within a shorter period of neurogenesis than the neocortical counterpart. Interestingly, many of the *Neurog1^CreER^* MADM clones populated 2-3 different thalamic nuclei, demonstrating that each round of radial glia division provides a pool of neurons that share similar birthdates and sometimes populate multiple nuclei. This is an efficient strategy to produce a large variety of neuronal types with a limited number of radial glial cells in a short period of time (summary schematics in Fig. S6B).

We further took advantage of the dual-color labeling of the progeny of IPCs and determined whether the terminal division of these cells produced neurons in the same or different thalamic nuclei. Although 78% of the final divisions resulted in neurons in the same nucleus, the remaining 22% produced neurons in two different nuclei. The most frequent pair of these nuclei was VP and dLG, the two principal sensory nuclei that share patterns of gene expression (Frangeul et al., 2016; Gezelius et al., 2016; Nagalski et al., 2016) and functional interactions (Moreno-Juan et al., 2017). Thus, the last division of thalamic IPCs may not be strictly symmetric, producing two neurons that have different nuclear fates. Alternatively, the division could be symmetric but post-mitotic mechanisms determine the fate of the two daughter neurons. Further studies will be needed to distinguish these possibilities.

## Materials and Methods

### Animals

*Gli1^CreERT2^* (Ahn and Joyner, 2005), *Olig3^CreERT2^* (Storm et al., 2009) and *Neurog1^CreERT2^* (Koundakjian et al., 2007; Bluske et al., 2012) mutants were found to provide an optimal labeling of thalamic progenitor cells at a clonal level when combined with the MADM-11 system (Hippenmeyer et al., 2010). We bred *Gli1^CreERT2/+^; MADM-11^GT/GT^* mice, *Olig3^CreERT2/+^; MADM-11^GT/GT^* mice or *Neurog1^CreERT2/+^; MADM-11^GT/GT^* mice with *MADM-11^TG/TG^* mice. A single dose of tamoxifen (132mg/kg body weight for *Gli1^CreERT2^,* 24mg/kg or 34mg/kg for *Olig3^CreERT2^* and *Neurog1^CreERT2^)* was administered to pregnant females (intraperitoneal injection for *Gli1^CreERT2^* and oral gavage for *Olig3^CreERT2^* and *Neurog1^CreERT2^)* at various time points (embryonic day 9.5 (E9.5) for *Gli1^CreERT2^,* E10.5 or E11.5 for *Olig3^CreERT2^* and E11.5 to E14.5 for *Neurog1^CreERT2^).* Embryos or postnatal pups were fixed and frozen. Each Cre driver line was also bred with *H2B-GFP* (for *Gli1^CreERT2/+^)* or Ai6 ZSGreen reporter (for *Olig3^CreERT2^* and *Neurog1^CreERT2/+^)* mice for a population-based lineage tracing with tamoxifen administration (60mg/kg body weight for *Olig3^CreERT2^* and *Neurog1^CreERT2/+^,* 132mg/kg body weight for for *Gli1^CreERT2/+^).* For the birth-dating study, ethynyldeoxyuridine (EdU; Invitrogen or Carbosynth) was injected intraperitoneally (50mg/kg body weight) into pregnant CD1 female mice at E9.5, 10.5, E11.5, E12.5, E13.5 or E14.5 and pups were perfusion fixed at on the day of birth (P0). All animal procedures used in this study were performed in accordance with the protocol approved by the Institutional Animal Care and Use Committee of Johns Hopkins University School of Medicine and University of Minnesota.

### Immunostaining, EdU staining, Confocal Imaging, and 3D reconstruction

Serial coronal brain sections (40μm in thickness) through the entire forebrain were cut by a cryostat or a sliding microtome, and were immunostained with the following antibodies: anti-GFP (1:200 or 500; goat; Rockland), anti-GFP (1:500; chicken; Aves Labs), anti-RFP (1:200 or 1,000; rabbit; Rockland; to detect tdTomato), anti-Nestin (1:500; chicken; Aves Labs), anti-goat/chicken Cy2, anti-rabbit Cy3 and anti-goat Cy5 (1:200 or 500; donkey; Jackson ImmunoResearch). EdU was visualized on cryosections in the detection solution (5uM Sulfo-Cy3 azide (Lumiprobe), 0.1M Tris pH7.5, 4mM copper sulfate, 100mM sodium ascorbate) for 30min after permeabilization in 0.5% Triton-X100 for 30min. Consecutive sections covering individual clones were imaged using Zeiss LSM 710 confocal microscope (Carl Zeiss). For 3D reconstruction, optical stacks from the entire diencephalon were serially aligned along the rostro-caudal axis using Reconstruct 1.1.0 (J.C. Fiala, NIH), followed by import into Imaris (Bitplane) for further analysis.

### Axial nomenclature and identification of thalamic nuclei

We adopted the axial nomenclature used in the prosomeric model of forebrain organization ((Puelles et al., 2013; Vue et al., 2007); Fig. 1A). In this system, the rostral border of the thalamus is the zona limitans intrathalamica (ZLI), and in the frontal view of cross sections, the bottom part (closer to the ZLI) is ventral-rostral and the top part (closer to the pretectum) is dorsal-caudal. Axial nomenclature of E18.5 sections (e.g., Fig. 6A) also follows this rule in order to retain consistency. For identification of thalamic nuclei within tissue sections, we referred to “Atlas of the Developing Mouse Brain at E17.5, P0 and P6” (Academic Press, 2006) and Nakagawa and O'Leary, (2001). Furthermore, we performed *in situ* hybridization (Vue et al., 2007) on E18.5 and P21 sections of wild type brains using the same orientation and thickness as used for the MADM brains, and generated a custom atlas (Fig. S4A,B). The mRNA probes for *Gbx2, Calb2, RORα, Foxp2, Igsf21, Slc6a4* and *Zyx* were used to label thalamic nuclei. At E18.5, *Gbx2* was expressed in PV, MD, CL, CM, Re/Rh, Po, LP, VM, MGv and the “medial ventral field” (shown by a white asterisk), which is not labeled in the available atlas (Paxinos, 2006) or book (Jones, 2007). Because this region is marked by ZSGreen in *Olig3^CreERT2^:* Ai6 reporter mice at E18.5 (Fig. S4C), we consider it to be thalamus-derived and included it in the analysis; *Calb2* was expressed in PV, LD, CL, CM, Re/Rh, Po, LP and medial ventral field; *RORα* was expressed in AD, MD, dLG, VP, MGv; *Foxp2* was expressed in PV, LD, CM, Re/Rh, MD, Po, LP, PF; *Isgf21* was expressed in AD, AV, AM, PV, LD, MD, Po, LP, VP, dLG, MGv, PF; *Slc6a4* was expressed in VP, dLG, MGv; *Zyx* was expressed in AV, PV, LD, MD, VL, VM, CM, Re/Rh, Po, PF. At P21, expression patterns remained essentially unchanged from E18.5 for all markers except *RORα*, which was expressed in an additional set of nuclei including PT, AD, AV, AM, LD, VL, VM, Po, LP and MGd.

### Computational Quantitative Analyses

Each clone was first assigned a vector of values representing its cellular contribution to each nucleus *(nucleus X, number of labeled cells in nucleus* X). Analogous to analysis of large datasets of gene expression in single-cell RNA sequencing in which each cell has a corresponding vector of values based on relative expression levels for each gene *(gene X, expression level of gene* X), we performed logarithmic transformation, followed by principal component analysis and unsupervised hierarchical clustering analysis. For principal component analysis, the clonal distribution data were rescaled such that the average number of cells in each nucleus was centered to zero with principal components as normalized eigenvectors of the covariance matrix of the cellular distributions. The clones were then ordered according to their contribution to the variance in the dataset. For unsupervised hierarchical clustering analysis, we calculated Euclidean distance among clones for plotting dendrograms and heatmaps. Since there were no negative values in the dataset, the Canberra distance was further analyzed for hierarchical clustering with Ward's linkage, which yielded similar results.

### Statistical Analyses

Computational analysis of MADM clones was performed using Excel and R software. GraphPad Prism ver.5.0 was used for other statistical tests. In Fig. 4C, statistical significance was assessed by two-tailed unpaired Student’s t-tests. *\*P* < 0.05; \*\**P* < 0.01; \*\*\**P* < 0.001. For the non-normal data, statistical significance was evaluated by Mann Whitney test. \**P* < 0.05; \*\**P* < 0.01.

## Acknowledgments

We thank Carmen Birchmeier for providing *Olig3^CreERT2^* mice, Jane Johnson and Lisa Goodrich for providing *Neurog1^CreERT2^* mice, Z. Josh Huang for providing *Rosa26- H2B-GFP* mice, Kimberly Christian, Steven McLoon, Timothy Monko, Bee Hua Chua, Ming Hui Ong and members of Ming and Song laboratories for comments on the manuscript, Rachel Larsen, Carmen Tso, Thomas Bao, Morgan McCullough, Shaylene McCue, Lihong Liu and Yuan Cai for technical support. The work was supported by grants from National Institutes of Health HD086820 to H.S and Y.N., R37NS047344 and P01NS097206 to H.S., MH110160, MH105128 and NS097370 to G.L.M., grants from National Natural Science Foundation of China (31771131) and Strategic Priority Research Program of the Chinese Academy of Sciences (XDA16020100) to Q.F.W and Undergraduate Research Opportunities Program (UROP) of University of Minnesota to E.P.S, E.B and M.F.

## Figure Legends

**Figure S1.**
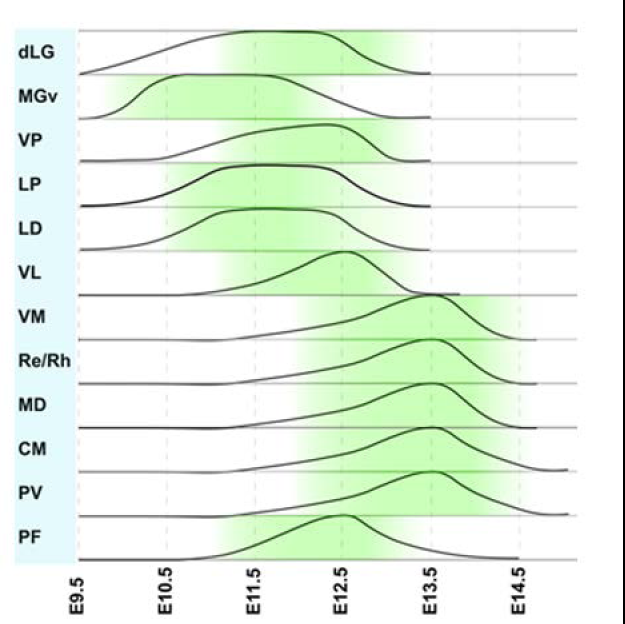
A qualitative schematic summary of EdU birthdating analysis shown in Figure 1B. Embryonic stages at the bottom shows the stage of EdU administration. dLG; dorsal lateral geniculate, MGv; ventral medial geniculate, VP; ventral posterior, LP; lateral posterior, LD; laterodorsal, VL; ventrolateral, VM; ventromedial, Re/Rh, reuniens/rhomboid, MD; mediodorsal, CM; centromedian, PV; paraventricular, PF; parafascicular.

**Figure S2.**
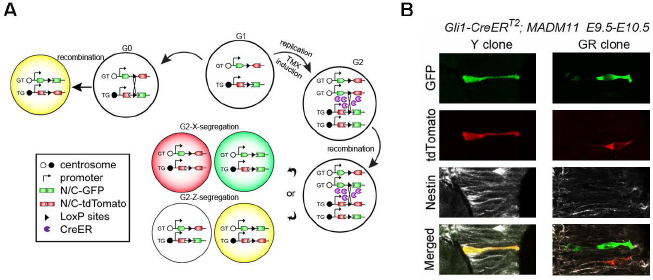
(**A**) In the MADM system, tamoxifen (TMX)-inducible Cre mice are used to drive interchromosomal recombination of the gene locus that contains partial coding sequences of EGFP and tdTomato (MADM-11GT and MADM-11TG). The recombination causes the production of fluorescent EGFP or tdTomato proteins. When the recombination occurs in a progenitor cell during G2 phase of the cell cycle, the two daughter cells express different combinations of fluorescent proteins depending on the mode of segregation of sister chromatids. Upon “G2-X” type segregation, one daughter cell expresses EGFP (green or “G” cell) and the other expresses tdTomato (red or “R” cell). If these daughter cells undergo further divisions, respective fluorescent proteins continue to be expressed in the progeny, generating a “G/R clone” that contains a mixture of green and red cells. The other, “G2-Z” type segregation produces one daughter cell that expresses both EGFP and tdTomato (yellow or “Y” cell) and the other daughter cell that expresses neither of the proteins. Thus, the G2-Z segregation generates a “Y clone”, which allows us to trace only half of the progeny of the recombined progenitor cell. (**B**) Immunostaining of an E10.5 section of *Gli1^CreERT2^* labeled clones with anti-Nestin antibody. Tamoxifen was administered at E9.5.

**Figure S3.**
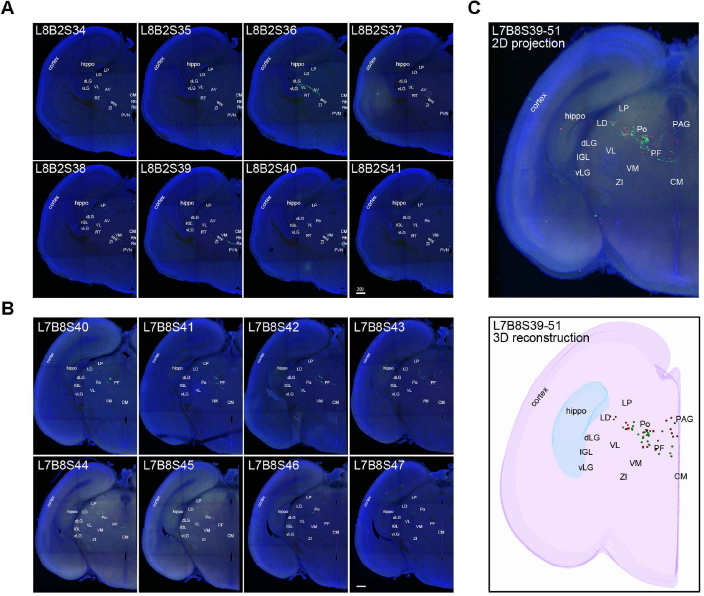
**(A-B)** Sample serial confocal images from two E18.5 *Gli1* MADM clones (L8B2 and L7B8). Each image is a confocal Z-stack of an individual section (40μm-thick) and shows the distribution of green and red cells in multiple thalamic nuclei. L8B2S34 to L8B2S41 as well as L7B8S40 to L7B8S47 represent eight consecutive sections. Scale bar: 200μm. (**C**) 2D projection and 3D reconstruction of an entire E18.5 *Gli1* MADM clone (L7B8) encompassing 13 sections. This clone lacks a retained RGC as shown.

**Figure S4.**
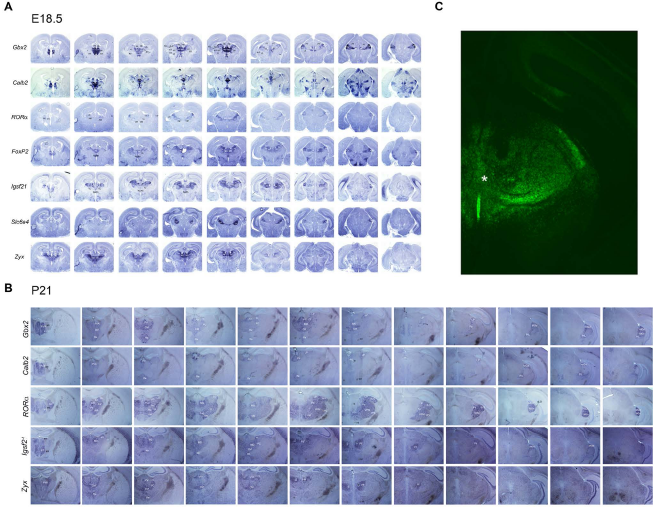
A custom atlas generated by *in situ* hybridization to define individual thalamic nuclei at E18.5 (**A**) and P21 (**B**). Expression of seven representative markers is shown. This is a consecutive set of 40μm-thick frontal sections. The left column is most dorsal and the right column is the most ventral (see Fig. 1A for axial orientation within the thalamus). Scale bar: 1mm. (**C**) An image of frontal section from *Olig3^CreERT2^;ZSGreen* brain to show labeling of medial ventral field (asterisk) by ZSGreen. Tamoxifen was administered at E9.5.

**Figure S5.**
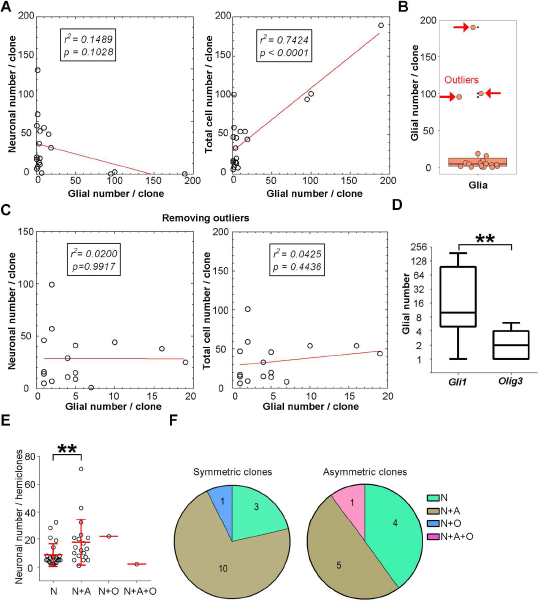
Characterization of glia-containing clones in the thalamus. (A) Scatter plots illustrating the relationship between the number of glia and neurons (left) or total cell number (right) in the P21 clones derived from both *Gli1* and *Olig3* brains. While the correlation between glial and neuronal number is not striking, the glial number is linearly correlated with total cell number in the postnatal clones. r^2^, linear correlation coefficient; we refer to statistically significant as p < 0.05. (**B**) The box plot overlaid with dot plot showing the distribution of glial number per clone. The red arrows indicate the outlier clones beyond the Gaussian distribution. These clones consisted mostly of glial cells. (**C**) The linear correlation analysis shows that there is no significant correlation between glial and neuronal number (left) or glial and total cell number (right) after removing the outlier clones from P21 brains. (**D**) The box plot showing the glial cell number in the *Gli1* and *Olig3* clones from P21 brains. **, p < 0.01 (Mann Whitney test). (**E**) Dot plot displaying the neuronal number in the hemi-clones that contain neurons only (N), neurons and astrocytes (N+A), neurons and oligodendrocytes (N+O) or neurons, astrocytes and oligodendrocytes (N+A+O). Each dot represents one hemi-clone and the red lines represent mean ± SEM. **, p < 0.01 (Mann Whitney test). (**F**) Pie chart showing the percentage of symmetric proliferative and asymmetric neurogenic clones that contain N, N+A, N+O or N+A+O.

**Figure S6.**
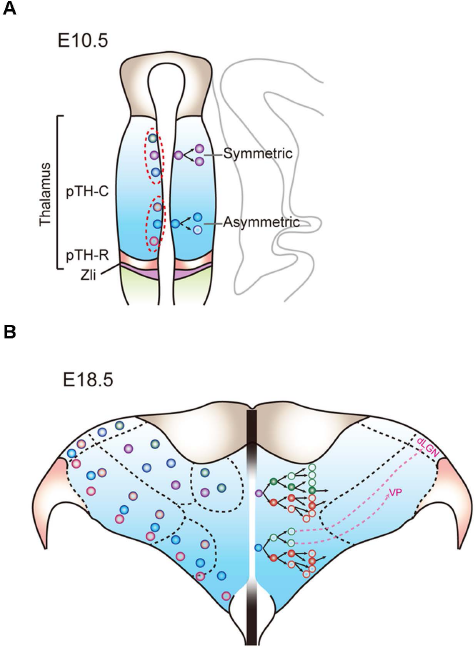
A schematic summary of the ontogenetic organization of thalamic nuclei. A schematic summary of the current study showing the main principles underlying spatiotemporal regulation of thalamic progenitor cell specification at E10.5 (**A**) and E18.5 (**B**). By E10.5, progenitor cells in the rostral-ventral part of the pTH-C domain are already undergoing asymmetric divisions (**A**, dots in the lower part of the pTH-C domain on the right side; see also Fig. 1A) and produce neurons that later populate principal sensory nuclei including ventral posterior (VP) and dorsal lateral geniculate (dLG). (**B**) The long-term lineage tracing shows that within the cell lineages that are derived from rostral-ventral progenitor cells, earlier-born neurons populate laterally located, principal sensory nuclei including dLG and VP, whereas later-born neurons populate more medial nuclei (dots in the lower part of the thalamus in **B**). In contrast, progenitor cells at more caudo-dorsal locations are still mainly undergoing symmetric division at E10.5 (**A**, dots in the upper part of the pTH-C domain on the right side), and eventually produce neurons in caudo-dorsally located nuclei (dots in the upper part of the thalamus in **B**). Regardless of cell positioning, a majority of radial glial precursors undergo either symmetric proliferative or asymmetric neurogenic division in the first round of cell division after genetic labelling at E10.5 (see also Fig. 3D). On the left side of schematics, progenitor cells (**A**) and their progeny (**B**) are color-coded to indicate their lineage relationship.

## References

Ahn, S., and Joyner, A.L. (2005). In vivo analysis of quiescent adult neural stem cells responding to Sonic hedgehog. Nature 437, 894–897.

Altman, J., and Bayer, S.A. (1988). Development of the rat thalamus: I. Mosaic organization of the thalamic neuroepithelium. The Journal of comparative neurology 275, 346–377.

Angevine, J.B., Jr. (1970). Time of neuron origin in the diencephalon of the mouse. An autoradiographic study. The Journal of comparative neurology 139, 129–187.

Bluske, K.K., Vue, T.Y., Kawakami, Y., Taketo, M.M., Yoshikawa, K., Johnson, J.E., and Nakagawa, Y. (2012). β-Catenin signaling specifies progenitor cell identity in parallel with Shh signaling in the developing mammalian thalamus. Development 139, 2692–2702.

Bonaguidi, M.A., Wheeler, M., Shapiro, J.S., Stadel, R., Sun, G.J., Ming, G-l., and Song, H. (2011). In vivo clonal analysis reveals self-renew and multipotent adult neural stem cell characteristics. Cell 145, 1142–55.

Clasca, F., Rubio-Garrido, P., and Jabaudon, D. (2012). Unveiling the diversity of thalamocortical neuron subtypes. The European journal of neuroscience 35, 1524–1532.

Delogu, A., Sellers, K., Zagoraiou, L., Bocianowska-Zbrog, A., Mandal, S., Guimera, J., Rubenstein, J.L., Sugden, D., Jessell, T., and Lumsden, A. (2012). Subcortical visual shell nuclei targeted by ipRGCs develop from a Sox14+-GABAergic progenitor and require Sox14 to regulate daily activity rhythms. Neuron 75, 648–662.

Florio, M., Huttner, W.B. (2014). Neural progenitors, neurogenesis and the evolution of the neocortex. Development 141, 2182–2194.

Frangeul, L., Pouchelon, G., Telley, L., Lefort, S., Luscher, C., and Jabaudon, D. (2016). A cross-modal genetic framework for the development and plasticity of sensory pathways. Nature 538, 96–98.

Gao, P., Postiglione, M.P., Krieger, T.G., Hernandez, L., Wang, C., Han, Z., Streicher, C., Papusheva, E., Insolera, R., Chugh, K., et al. (2014). Deterministic progenitor behavior and unitary production of neurons in the neocortex. Cell 159, 775–788.

Garcia-Lopez, R., Vieira, C., Echevarria, D., and Martinez, S. (2004). Fate map of the diencephalon and the zona limitans at the 10-somites stage in chick embryos. Dev Biol 268, 514–530.

Gezelius, H., Moreno-Juan, V., Mezzera, C., Thakurela, S., Rodriguez-Malmierca, L.M., Pistolic, J., Benes, V., Tiwari, V.K., and Lopez-Bendito, G. (2016). Genetic Labeling of Nuclei-Specific Thalamocortical Neurons Reveals Putative Sensory-Modality Specific Genes. Cerebral cortex (New York, NY: 1991).

Hippenmeyer, S., Youn, Y.H., Moon, H.M., Miyamichi, K., Zong, H., Wynshaw-Boris, A., and Luo, L. (2010). Genetic mosaic dissection of Lis1 and Ndel1 in neuronal migration. Neuron 68, 695–709.

Jager, P., Ye, Z., Yu, X., Zagoraiou, L., Prekop, H.T., Partanen, J., Jessell, T.M., Wisden, W., Brickley, S.G., and Delogu, A. (2016). Tectal-derived interneurons contribute to phasic and tonic inhibition in the visual thalamus. Nature communications 7, 13579.

Jeong, Y., Dolson, D.K., Waclaw, R.R., Matise, M.P., Sussel, L., Campbell, K., Kaestner, K.H., and Epstein, D.J. (2011). Spatial and temporal requirements for sonic hedgehog in the regulation of thalamic interneuron identity. Development (Cambridge, England) 138, 531–541.

Jones, E.G. (2007). The Thalamus. Cambridge University Press.

Kohwi, M., and Doe, C.Q. (2013). Temporal fate specification and neural progenitor competence during development. Nat Rev Neurosci 14, 823–838.

Koundakjian, E.J., Appler, J.L., and Goodrich, L.V. (2007). Auditory neurons make stereotyped wiring decisions before maturation of their targets. The Journal of neuroscience: the official journal of the Society for Neuroscience 27, 14078–14088.

Kriegstein, A., Alvarez-Buylla, A. (2009). The glial nature of embryonic and adult neural stem cells. Annual review of neuroscience 32, 149–184.

Mihalas, A.B., Elsen, G.E., Bedogni, F., Daza, R.A.M., Ramos-Laguna, K.A., Arnold, S.J., Heyner, R.F. (2016) Intermediate progenitor cohorts differentially generate cortical layers and require Tbr2 for timely acquisition of neuronal subtype identity. Cell reports 16, 92–105.

Miyata, T., Kawaguchi, A., Okano, H., and Ogawa, M. (2001). Asymmetric inheritance of radial glial fibers by cortical neurons. Neuron 31, 727–741.

Moreno-Juan, V., Filipchuk, A., Anton-Bolanos, N., Mezzera, C., Gezelius, H., Andres, B., Rodriguez-Malmierca, L., Susin, R., Schaad, O., Iwasato, T., et al. (2017). Prenatal thalamic waves regulate cortical area size prior to sensory processing. Nature communications 8, 14172.

Nagalski, A., Puelles, L., Dabrowski, M., Wegierski, T., Kuznicki, J., and Wisniewska, M.B. (2016). Molecular anatomy of the thalamic complex and the underlying transcription factors. Brain structure & function 221, 2493–2510.

Nakagawa, Y., and O'Leary, D.D. (2001). Combinatorial expression patterns of LIM-homeodomain and other regulatory genes parcellate developing thalamus. The Journal of neuroscience: the official journal of the Society for Neuroscience 21, 2711–2725.

Paxinos, G. (2006). Atlas of the Developing Mouse Brain at E17.5, P0 and P6 (Academic Press).

Petrof, I., and Sherman, S.M. (2009). Synaptic properties of the mammillary and cortical afferents to the anterodorsal thalamic nucleus in the mouse. Journal of neuroscience 29, 7815–7819.

Puelles, L., Harrison, M., Paxinos, G., and Watson, C. (2013). A developmental ontology for the mammalian brain based on the prosomeric model. Trends in neurosciences 36, 570–578.

Sherman, S.M. (2016). Thalamus plays a central role in ongoing cortical functioning. Nature neuroscience 19, 533–541.

Sherman, S.M., and Guillery, R.W. (2013). Thalamocortical Processing: Understanding the Messages that Link the Cortex to the World (MIT Press).

Shi, W., Xianyu, A., Han, Z., Tang, X., Li, Z., Zhong, H., Mao, T., Huang, K., and Shi, S.H. (2017). Ontogenetic establishment of order-specific nuclear organization in the mammalian thalamus. Nature neuroscience 20, 516–528.

Storm, R., Cholewa-Waclaw, J., Reuter, K., Brohl, D., Sieber, M., Treier, M., Muller, T., and Birchmeier, C. (2009). The bHLH transcription factor Olig3 marks the dorsal neuroepithelium of the hindbrain and is essential for the development of brainstem nuclei. Development (Cambridge, England) 136, 295–305.

Suzuki-Hirano, A., Ogawa, M., Kataoka, A., Yoshida, A.C., Itoh, D., Ueno, M., Blackshaw, S., and Shimogori, T. (2011). Dynamic spatiotemporal gene expression in embryonic mouse thalamus. The Journal of comparative neurology 519, 528–543.

Vue, T.Y., Aaker, J., Taniguchi, A., Kazemzadeh, C., Skidmore, J.M., Martin, D.M., Martin, J.F., Treier, M., and Nakagawa, Y. (2007). Characterization of progenitor domains in the developing mouse thalamus. The Journal of comparative neurology 505, 73–91.

Vue, T.Y., Bluske, K., Alishahi, A., Yang, L.L., Koyano-Nakagawa, N., Novitch, B., and Nakagawa, Y. (2009). Sonic hedgehog signaling controls thalamic progenitor identity and nuclei specification in mice. The Journal of neuroscience: the official journal of the Society for Neuroscience 29, 4484–4497.

Wang, L., Bluske, K.K., Dickel, L.K., and Nakagawa, Y. (2011). Basal progenitor cells in the embryonic mouse thalamus - their molecular characterization and the role of neurogenins and Pax6. Neural development 6, 35.

Yao B, Christian KM, He C, Jin P, Ming GL, Song H. (2016). Epigenetic mechanisms in neurogenesis. Nature Review Neuroscience 17, 537–49.

Zong, H., Espinosa, J.S., Su, H.H., Muzumdar, M.D., and Luo, L. (2005). Mosaic analysis with double markers in mice. Cell 121, 479–492.

